# Attentional capture by fearful faces requires consciousness and is modulated by task-relevancy: a dot-probe EEG study

**DOI:** 10.1101/2023.02.07.527584

**Authors:** Zeguo Qiu, Jiaqin Jiang, Stefanie I. Becker, Alan J. Pegna

**Affiliations:** School of Psychology, The University of Queensland, Brisbane 4072, Australia

**Keywords:** spatial attention, awareness, fearful faces, EEG, mass univariate analysis, multivariate pattern analysis

## Abstract

In the current EEG study, we used a dot-probe task in conjunction with backward masking to examine the neural activity underlying awareness and spatial processing of fearful faces and the neural processes for subsequent cued spatial targets. We presented face images under different viewing conditions (subliminal and supraliminal) and manipulated the relation between a fearful face in the pair and a subsequent target. Through both mass univariate analysis and multivariate pattern analysis, we found that fearful faces can be processed to an extent where they attract spatial attention only when they are presented supraliminally and when they are task-relevant. The spatial attention capture by fearful faces also modulated the processing of subsequent lateralised targets that were spatially congruent with the fearful face, in both behavioural and neural data. There was no evidence for nonconscious processing of the fearful faces in the current paradigm. We conclude that spatial attentional capture by fearful faces requires visual awareness and it is modulated by top-down task demands.

## 1. Introduction

Fearful expressions communicate information to other individuals regarding our perception of the environment. Specifically, we may express fear in response to dangerous events or threat. Therefore, fearful faces are usually perceived as indicators of negative events and they tend to attract our attention easily (Pourtois et al., 2004; Schupp et al., 2004). It has been reported that emotional faces including fearful faces can be detected faster and they elicit stronger neural activity, compared to neutral faces (for a review see Schindler & Bublatzky, 2020), even when people are unaware of them (Qiu et al., 2022c; Tamietto & De Gelder, 2010; Vuilleumier, 2005).

With regards to attentional capture, it has been shown that the presence of a fearful face can enhance the processing of a subsequent stimulus. Such modulatory effects of fearful faces on subsequent targets have been mainly examined using the dot-probe paradigm (e.g., Torrence & Troup, 2018). In this paradigm, a pair of face stimuli is presented before a lateralised target stimulus. The lateralised target can be presented in the same spatial location as the emotional face that precedes it (the congruent condition), or at the location opposite to the emotional face (the incongruent condition). The response to the lateralised target is measured, and the differences between congruent and incongruent conditions can be used as an index of spatial attention to the preceding faces. Previous research has repeatedly shown that a target (e.g., a dot or a letter) can be detected faster (Carlson & Reinke, 2008, 2010; Torrence et al., 2017) or discriminated more accurately (Pourtois et al., 2004) when it follows the emotional face in the spatially congruent condition, compared to when it is spatially incongruent to the emotional face (but see van Rooijen et al., 2017).

Neural imaging studies have provided supporting evidence for these behavioural observations. For example, in a dot-probe experiment using electroencephalography (EEG) recording, Carlson and Reinke (2010) presented participants with pairs of faces as the cues and a lateralised dot as the target. At the behavioural level, they found that participants’ reaction time towards dots in the congruent condition was shorter than the incongruent condition. Additionally, the neural activity, indexed by event-related potentials (ERPs), at posterior electrodes were found to be enhanced by a lateralised fearful face compared to a neutral face. The magnitude of the increase in the ERPs, in particular the face-sensitive N170, positively correlated with the reaction time difference between congruent and incongruent trials. These results were taken to suggest that fearful faces attracted spatial attention and facilitated task performance for congruent targets. However, to show that fearful faces indeed modulate attention to the subsequent targets, it would have been more compelling to show that fearful faces alter the neural response to the subsequent targets. The target-related ERPs were not reported in the study (Carlson & Reinke, 2010), leaving it an open question how attention to fearful faces modulates neural activity for the targets.

In an object-substitution masking study by Giattino and colleagues (2018), face and house images were used as cueing stimuli, presented subliminally (for 17ms) and subsequently masked. Participants were required to detect a target rectangle that was either validly or invalidly cued. It was found that the participants’ ability to detect the cue stimuli was no different from chance-level guessing (Giattino et al., 2018). However, participants’ early neural activity (i.e., P1) in response to the target stimuli was enhanced when the targets were validly cued, even in trials where participants reported not being aware of the preceding cue. This cue validity effect was also found in the behavioural data such that participants localised the targets faster and more accurately in the congruent condition, even when participants were not aware of the cues (Giattino et al., 2018). Thus, in the absence of awareness of face cues, the spatial information about them was processed to a level where it modulated the neural responses to the subsequent stimuli.

In a series of backward masking experiments, we analysed ERPs for fearful faces presented in face pairs with different visibility (subliminal and supraliminal viewing conditions; Qiu et al., 2022a, 2023). We used 16ms of presentation time in the subliminal viewing condition, as opposed to 33ms which was used in Carlson and Reinke (2010), for a stronger impeding effect on visual awareness (Milders et al., 2008). Our results showed that fearful faces can attract spatial attention by eliciting an N2-posterior-contralateral (N2pc), only when the faces were presented above the awareness threshold (266ms; supraliminal viewing) and when they were relevant to participants’ tasks (Qiu et al., 2022a). Although subliminal fearful faces did not elicit an N2pc, some fear-related non-spatial enhancement effect is present in the data (Qiu et al., 2022d). We then ask whether any of the fear-related effects are sustained and can modulate the neural response to stimuli presented after the faces (i.e., modulate target-related EEG signals). Importantly, we ask whether any of the effects require visual awareness, or they can occur as nonconscious processes.

To answer these questions, in the current study, we used a dot-probe task together with the backward masking technique. Specifically, pairs of faces were presented either briefly (for 16ms) or for a longer time (166ms) and immediately backward masked. A following lateralised dot either appeared on the same side as the fearful face (congruent) or on the side opposite to it (incongruent). In one half of the experiment, the participants were required to respond to the target dots as well as the faces that preceded them, whereas in the other half of the experiment, participants were instructed to ignore the faces.

For the face stimuli, we expected to find an N2pc for the fearful face only in the supraliminal viewing condition, replicating our previous finding (Qiu et al., 2022a). For the target dot stimuli, we expected dots in the congruent conditions to be detected faster than the incongruent condition. We predicted that the early ERPs (e.g., P1) for congruent dots would be enhanced compared to incongruent dots. Further, if fearful faces can be processed nonconsciously, such cue validity effect should be observed in both supraliminal and subliminal face presentations. Data were also analysed with a multivariate approach to examine neural patterns associated with the variables of interest (i.e., fearful face location, congruency) which may not be revealed in univariate ERP analyses.

## 2. Materials and method

### 2.1 Participants

We determined the sample size in MorePower (Campbell & Thompson, 2012) using an effect size from a previous study with a similar design (*η*_*p*_^*2*^ = 0.22; Carlson & Reinke, 2008). A minimum of 24 participants were required for a significant main effect of congruency in reaction time in a 3(fearful-face-dot congruency: congruent, incongruent, control) x 2(face-visibility: subliminal, supraliminal) x 2(face-relevancy: relevant, irrelevant) design with an effect size of 0.22 (power = 0.9, two-tailed alpha = 0.05). Thirty-one participants were recruited and were compensated with either course credits or $40 AUD. Data from five participants were excluded after data pre-processing (see below). Therefore, 26 participants constituted the final sample (*M*_*age*_ = 21.9, *SD*_*age*_ = 2.1, 8 males, 18 females). This study was approved by the University of Queensland ethics committee.

### 2.2 Apparatus and experimental stimuli

The experiment was programmed and run in PsychoPy 3 (Peirce et al., 2019) and all stimuli were presented on a 24-inch ASUS LCD monitor (resolution: 1920 × 1080 pixels) placed 70 cm away from the participant’s eyes.

Face stimuli were obtained from the Radboud Face Database (Langner et al., 2010). We used fearful and neutral face images from 10 different models (five females and five males). Non-face information including hair was removed by cutting the face images into oval shapes (6.5° × 5.1° in visual angle; see Figure 1A). The mask stimuli were created by scrambling the neutral face images for each model using the Scramble Filter tool (http://telegraphics.com.au/sw/product/scramble) such that each mask image consisted of 208 randomly scrambled squares (4.4 mm × 4.4 mm each), see Figure 1B. In this experiment, we used a bilateral presentation of faces and mask stimuli. Each lateralised stimulus was presented 4.1° (in visual angle) away from a central fixation on the screen. The possible face combinations included a) fearful face on the left and neutral face on the right (fearful-face-on-left); b) neutral face on the left and fearful face on the right (fearful-face-on-right); c) two neutral faces.

**Figure 1.**
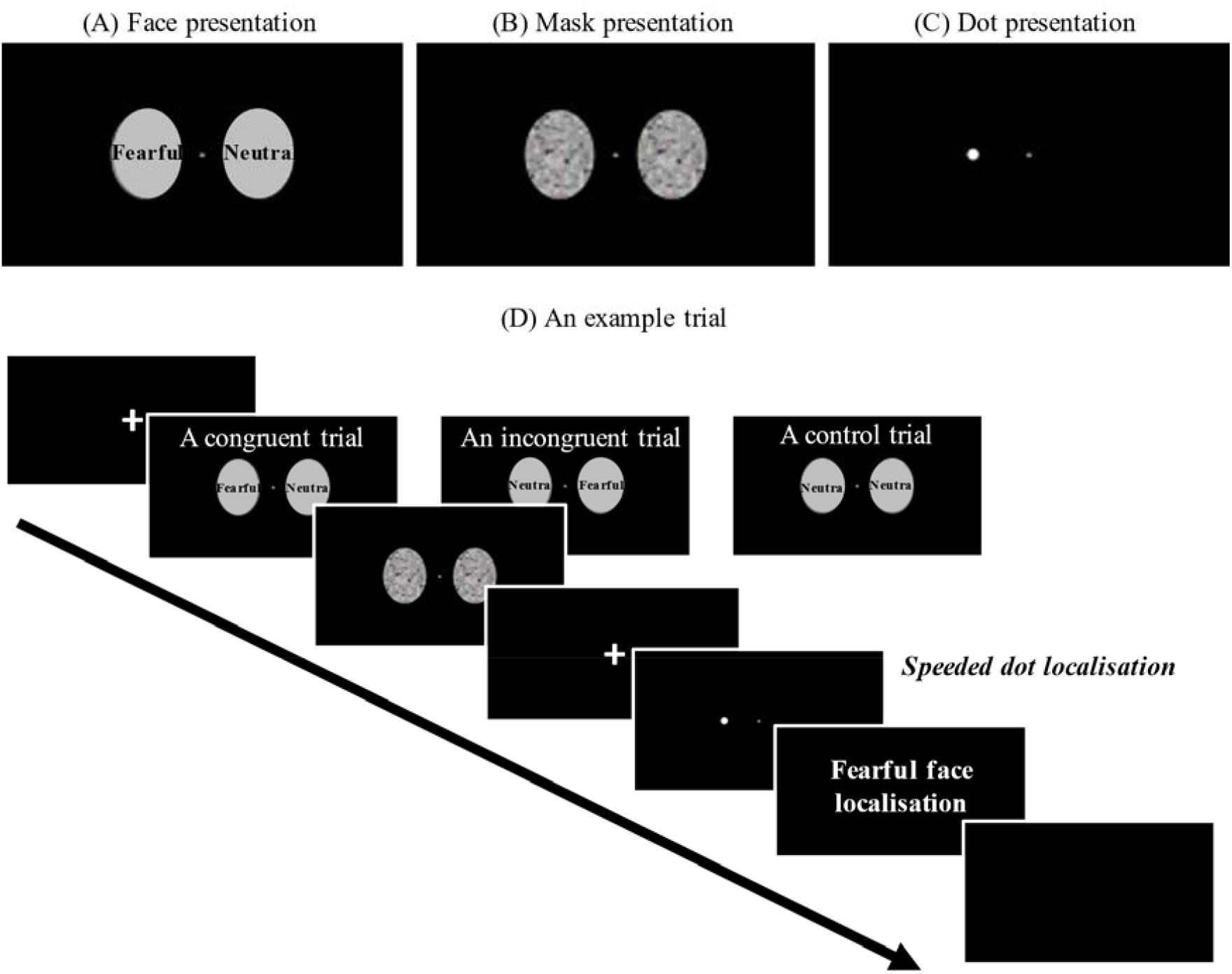
Examples of (A) the face images (Fearful-on-left) and (B) mask images. (C) An example of the lateralised target dot (Dot-on-left). (D) The full sequence of a trial. Note that the fearful face localisation task was only required in the face-relevant conditions. The face stimuli are covered and de-identified in this picture in accordance with bioRxiv policies. Actual face images were used in the experiment.

A white disc (a dot) extending 0.25° × 0.25° in visual angle was used as the target stimulus in the current dot-probe paradigm (Figure 1C). The distance between the centre of the dot and the central fixation was 4.1° in visual angle.

All images were rendered black-and-white and were presented on a black screen. Image editing was performed in Adobe Photoshop (version 22.4.0).

### 2.3 Procedure

As shown in Figure 1D, at the start of each trial, a fixation screen was presented with a variable duration between 500-800ms. Then, a pair of face stimuli appeared for either 16ms (subliminal) or 166ms (supraliminal), immediately followed by a pair of mask stimuli for either 166ms or 16ms, making the total duration of faces and masks the same across conditions. Afterwards, a fixation screen was presented for 66ms (Torrence et al., 2017) and was followed by a lateralised dot presented either on the left or the right side of the screen for 750ms. In a small proportion of the trials (360 trials in total), there was no dot following the mask (baseline condition). The baseline trials were introduced for us to obtain clean dot-related ERPs (see Data analysis). Upon the onset of the dot presentation (or the blank screen in the baseline condition), participants were required to correctly localise the dot as quickly as possible (left arrow key = dot on left; right arrow key = dot on right; both keys = no dot) with their right hand. If no response was made within 1000ms after the onset of the dot, a prompt “Too slow!” would be presented on the screen.

There were two types of blocks in the experiment: face-relevant and face-irrelevant blocks. In the face-relevant blocks, participants were instructed to first perform the dot localisation task. Then, they were required to indicate on which side of the screen they saw a fearful face (Q key = fearful face on left; E key = fearful face on right; W = no fearful face/two neutral faces) with their left hand. In the face-irrelevant blocks, participants were instructed to respond only to the dots and ignore the faces. That is, they only needed to perform the speeded dot localisation task in these blocks.

There were three conditions regarding the location of the fearful face and the lateralised dot in each trial: *congruent* condition, where the fearful face and the subsequent dot were presented on the same side of the screen, *incongruent* condition, where they were presented on different sides of the screen and the *control* condition where the two neutral faces were presented before the dot.

There were 16 blocks of 1200 trials (including 360 baseline trials) in total with short breaks provided between blocks. Participants completed eight face-relevant blocks in either the first half or the second half of the experiment and completed the eight face-irrelevant blocks in the other half of the experiment. This order was determined randomly by the experimental program for each participant.

### 2.4 EEG data recording and pre-processing

Raw continuous EEG was recorded at 1024 Hz using the BioSemi ActiveTwo system (Biosemi, Amsterdam, Netherlands). Sixty-four electrodes were placed according to the international 10–20 system location. Horizontal electrooculogram (EOG) was recorded with two bipolar electrodes. Vertical EOG was recorded with an external electrode placed below participants’ left eye. Recordings were referenced online to the CMS/DRL electrodes.

Pre-processing of the EEG data was performed with EEGLAB (Delorme & Makeig, 2004) and ERPLAB (Lopez-Calderon & Luck, 2014). We interpolated electrodes that produced noise throughout the experiment. Signals were re-sampled to 512 Hz offline, filtered from 0.1 to 30 Hz and notch-filtered at 50 Hz to remove line noise. All signals were then re-referenced to the average of all electrodes. Signals were segmented into epochs with a time window of 600 ms from the onset of the faces, using a pre-stimulus baseline (−100 to 0 ms). Independent component analysis was performed on the epoched data to identify and remove eye-blink and eye-movement components in the signals. After eye-related components were removed, epochs containing other artefacts were detected and removed on a trial-by-trial basis through visual inspection. Consequently, data from five participants were excluded for further analyses due to the limited number of epochs remaining (i.e., fewer than 40 trials for each condition of interest). On average, 91% epochs were kept for the remaining participants (*N* = 26).

### 2.5 Data analysis

At the behavioural level, we examined the reaction time data from the speeded dot localisation task with a repeated-measures ANOVA. We only used reaction time data from the task-correct trials and excluded datapoints identified as outliers from an outlier check (i.e., beyond 3^rd^ quartile + 1.5*interquartile) for each participant. We did not analyse the accuracy data for the dot localisation task because the accuracy was near ceiling (percent correct: *M* = 0.96, *SD* = 0.03). For the fearful face localisation task, we examined the accuracy data with a paired-samples *t*-test. All behavioural data analyses were performed in IBM SPSS Statistics 27.

For the EEG data, we separately examined the face-related signals and dot-related signals, using both a univariate approach and a multivariate method.

#### 2.5.1 Mass univariate analysis (ERP analysis)

We conducted ERP analyses using the factorial mass univariate analysis toolbox (Fields & Kuperberg, 2020) and the mass univariate analysis toolbox (for pairwise comparisons; Groppe et al., 2011).

Because we were interested in the early visual processing of the stimuli, posterior electrodes were selected as regions of interest. Only lateral electrodes were included in ERP analyses because we were interested in ERP components that are calculated as difference waves between lateral electrodes (i.e., N2pc). As a result, analyses were performed on electrodes P3/4, P5/6, P7/8, P9/10, PO3/4, PO7/8 and O1/2. For significant difference testing, we performed the cluster-based permutation test (10000 permutations) on all time-points within the epoch (0-600ms), with a family-wise α level of 0.05. Electrodes were considered as spatial neighbours if adjacent electrodes were within 3.3cm from each other (Mean spatial neighbours = 2.9; cluster inclusion *p* < .05). Follow-up comparisons were conducted using the cluster-based permutation *t*-tests (two-tailed family-wise α = .05).

For the analyses on the face-related signals, we collapsed the left and right hemispheres and retained the information about the relation between the location of the fearful face and hemispheres: face contralateral to the hemisphere and face ipsilateral to the hemisphere. For the neutral faces condition, the average between the left and right hemispheres were calculated.

For the analyses on the dot-related signals, we first subtracted signals in the dot-absent baseline trials from the dot-present trials to remove effects from the preceding face stimuli. Specifically, the ERPs time-locked to the dot onset from the baseline condition would be subtracted from the ERPs from the experimental condition (i.e., dot-present) that was preceded by the same face combination (e.g., fearful-on-left). Analyses were then performed on the baseline-subtracted ERPs.

#### 2.5.2 Multivariate pattern analysis (MVPA)

We also examined the data with a multivariate approach using the CoSMoMVPA toolbox (Oosterhof et al., 2016) and LIBSVM (Chang & Lin, 2011). We used a radial kernel support vector machine on each time point to find the decision boundary that discriminated between patterns of two conditions of interest using signals across 16 posterior electrodes (P3/4, P5/6, P7/8, P9/10, PO3/4, PO7/8, O1/2, POz and Oz). Then, data were spatially filtered with surface Laplacian (Kayser & Tenke, 2006) and temporally smoothed with a Gaussian-weighted running average of 20ms. Classification was performed on each time point and 4 neighbouring time points to avoid the overfitting issue (Grootswagers et al., 2017). Single-trial data were partitioned into 10 chunks and the two classification targets were equally likely to occur in each chunk. Following a leave-one-out procedure, each classifier at each time point was trained on data from nine chunks and tested on the remaining chunk. Decoding accuracies of all iterations were then averaged at each time point for each participant. Statistical significance testing was conducted using one-sample *t*-tests (against chance-level decoding performance at 50%). The *t* statistics were corrected for multiple comparisons using Threshold-Free Cluster Enhancement (TFCE) and Monte Carlo-based permutations (Oosterhof et al., 2016). Briefly, a null distribution was acquired through flipping the sign of the statistics across time points for a random half of participants, iteratively for 10000 times. The observed TFCE statistic at each time point was considered significant if its value was larger than the 95th percentile of the null distribution (i.e., *p* < .05 for a one-tailed test; Smith & Nichols, 2009).

For the face-related signals, we decoded the spatial location of the fearful face (fearful-face-on-left vs. fearful-face-on-right) for each condition of face-visibility (collapsing across face-relevancy), and decoded the fearful face location for each condition of face-relevancy (collapsing across face-visibility).

For the dot-related signals, we performed decoding of fearful-face-dot congruency (congruent vs. incongruent) across all conditions and also separately for the face-subliminal and face-supraliminal conditions.

## 3. Results

### 3.1 Behavioural data

#### 3.1.1 Fearful face localisation task

A paired-samples *t*-test on the accuracy data (percent correct) for the fearful face localisation task revealed that the fearful face was more accurately localised in the supraliminal condition (*M* = 0.74, *SD* = 0.19) than the subliminal condition (*M* = 0.34, *SD* = 0.02), *t*(25) = 10.51, *p* < .001, *d* = 2.06. A one-sample *t*-test showed that the accuracy in the subliminal condition was not different from chance-level performance (0.33), *t*(25) = 0.92, *p* = .368.

#### 3.1.2 Dot localisation task

A 3(fearful-face-dot congruency) × 2(face-visibility) x 2(face-relevancy) repeated-measures ANOVA on the reaction times (in seconds) revealed a significant main effect of face-visibility (*F*(1, 25) = 28.05, *p* < .001, *η*_*p*_^*2*^ = .53), a main effect of face-relevancy (*F*(1, 25) = 92.63, *p* < .001, *η*_*p*_^*2*^ = .79) and a significant interaction between the two, *F*(1, 25) = 27.01, *p* < .001, *η*_*p*_^*2*^ = .52. Specifically, participants were faster at localising the dot in the face-irrelevant condition (*M* = 0.34, *SD* = 0.04) than face-relevant condition (*M* = 0.44, *SD* = 0.06), and when the preceding faces were presented subliminally (*M* = 0.38, *SD* = 0.04) than when presented supraliminally (*M* = 0.40, *SD* = 0.05). The effect of face-relevancy was significant in both face-visibility conditions, *ps* < .001. However, the effect of face-visibility was significant only when the faces were task-relevant, *p* < .001. When participants did not need to attend to the faces, no difference was found between face-subliminal and face-supraliminal conditions, *p* = .295.

The interaction between face-visibility and congruency was also significant, *F*(2, 50) = 22.92, *p* = .008, *η*_*p*_^*2*^ = .20. Follow-up tests showed that, the effect of congruency was non-significant in the face-subliminal condition, *F*(2, 50) = 1.33, *p* = .274, but was significant in the face-supraliminal condition, *F*(2, 50) = 3.98, *p* = .025, *η*_*p*_^*2*^ = .14. As part of our planned comparisons, the levels of congruency were compared against each other in the face-supraliminal condition. We found that, the dot was localised significantly slower in the incongruent condition, compared to the congruent condition, *p* = .023, and the control condition, *p* = .018. The difference between congruent and control conditions was non-significant, *p* = .508. No other effect was significant, *Fs* < 2.25, *ps* >.116.

### 3.2 Mass univariate analysis

#### 3.2.1 Face-related ERPs

A 3(laterality based on the location of a fearful face: contralateral, ipsilateral, control) x 2(face-visibility: subliminal, supraliminal) × 2(face-relevancy: relevant, irrelevant) repeated-measures ANOVA revealed that all main effects were significant, *ps* < .05. As shown in Figure 2A, ERPs were significantly more negative in the supraliminal compared to the subliminal condition between 113-281ms (temporal peak: 215ms) with a maximal effect at P9/10, and between 293-594ms across all posterior electrodes. ERPs in the face-relevant condition were overall more negative than those in the face-irrelevant condition between 262-594ms (temporal peak: 348ms) with a maximal effect on P3/4, see Figure 2B. ERP waveforms are presented in Figure 2C.

**Figure 2.**
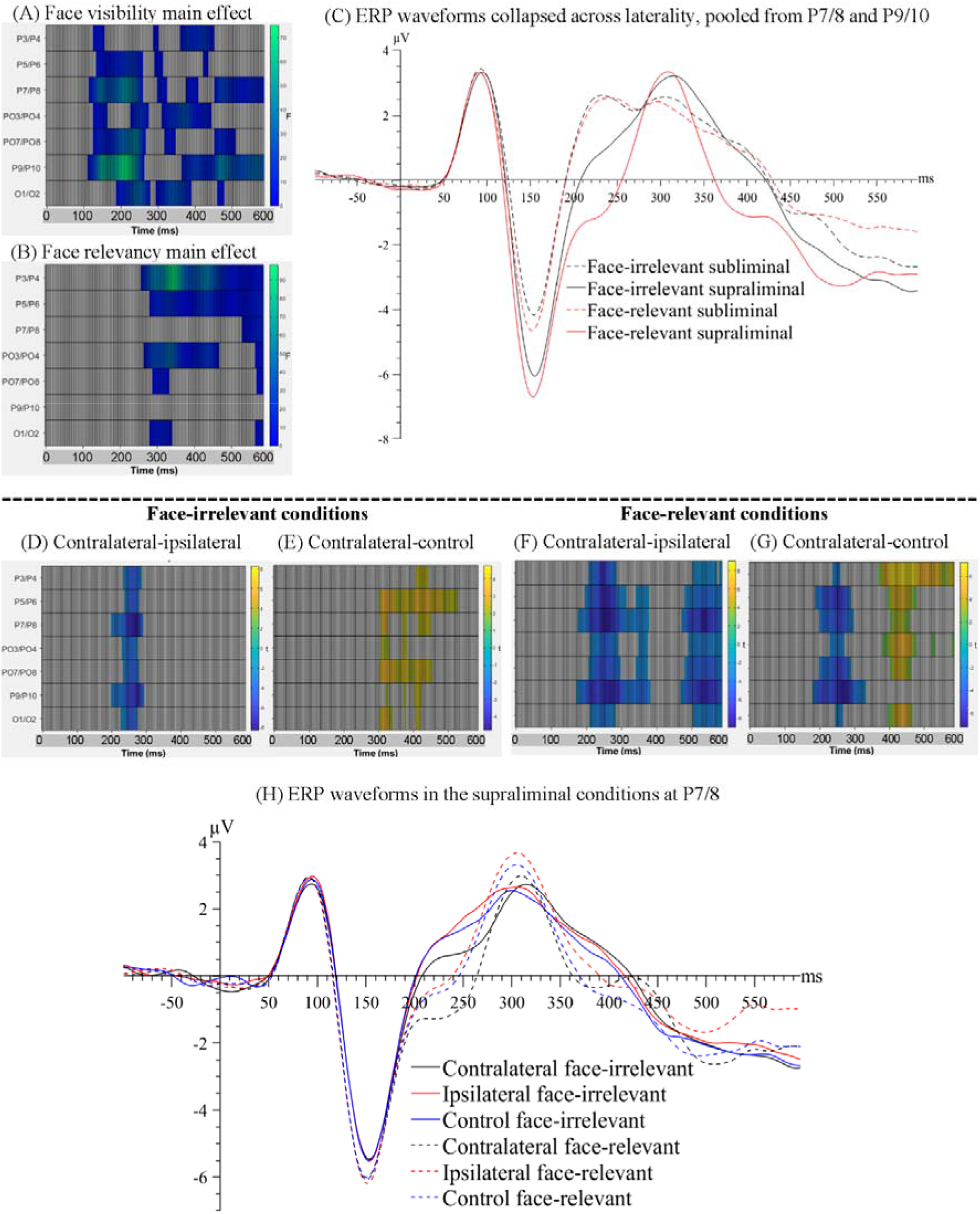
Raster plot of (A) the main effect of face-visibility and (B) the main effect of face-relevancy for face-related ERPs. (C) ERP waveforms collapsed across laterality conditions, pooled over P7/8 and P9/10, the two pairs of electrodes that showed the maximal main effect of face-visibility and the maximal interaction effect between face-visibility and face-relevancy. Raster plots for the contrasts between (D) contralateral and ipsilateral signals and between (E) contralateral and control signals, in the face-irrelevant conditions; between (F) contralateral and ipsilateral signals and between (G) contralateral and control signals, in the face-relevant conditions. (H) ERP waveforms for different conditions of laterality and face-relevancy in the supraliminal condition at electrodes P7/8, the pair that showed the maximal interaction effect between laterality and face-relevancy.

All interaction effects were significant including the three-way interaction between laterality, face-visibility and face-relevancy, *ps* < .05. Follow-up tests revealed that the effect of laterality was significant only in the supraliminal conditions. Thus, we compared three levels of laterality against each other in the supraliminal condition, separately for face-relevant and face-irrelevant conditions, using the cluster-based permutation *t*-tests.

When the supraliminally-presented faces were task-irrelevant, signals contralateral to the fearful face were more negative than ipsilateral signals between 203-297ms at posterior electrode sites (temporal peak: 273ms; spatial peak: P7/8; Figure 2D), reflecting an N2pc for the fearful face. Additionally, contralateral signals were more positive than signals in the control condition in a later time window spanning from 309 to 535ms (temporal peak: 332ms; spatial peak: PO7/8; Figure 2E).

When the supraliminally-presented faces were task-relevant, two negative clusters at posterior electrode sites were significant when comparing contralateral against ipsilateral signals: 176-387ms (temporal peak: 262ms; spatial peak: P7/8), again reflecting an N2pc for the target fearful face, and a later time window, 477-600ms (temporal peak: 543ms; spatial peak: P7/8), see Figure 2F. Contralateral signals were also more negative than signals in the control condition between 184-336ms (temporal peak: 273ms; spatial peak: P9/10; Figure 2G). The later negativity between 477-600ms in the contralateral-ipsilateral contrast (Figure 2F) likely reflected the sustained posterior contralateral negativity (SPCN; Luria et al., 2016), an ERP marker associated with working memory consolidation for task-relevant fearful faces. Note that the SPCN was not found in the face-irrelevant condition (Figure 2D).

One additional positive cluster was found when contrasting contralateral against control conditions: 402-484ms (temporal peak: 453ms; spatial peak: P3/4). Ipsilateral signals were also significantly more positive than control signals, as shown by a positive cluster spanning from 215 to 600ms (spatial peak: P3/4). It thus appears that the P300 was enhanced in the fearful-face-present condition (contralateral and ipsilateral conditions), compared to the control condition where both faces were neutral faces. The ERP waveforms are plotted in Figure 2H.

#### 3.2.2 Dot-related ERPs

To analyse on the dot-related ERPs, we averaged signals across the left and right electrode sites, and ran a 3(fearful-face-dot congruency) x 2(face-visibility) x 2(face-relevancy) repeated-measures ANOVA. As shown in Figure 3, the main effects of face visibility and face-relevancy, and the interaction between the two were significant, *ps* < .05.

**Figure 3.**
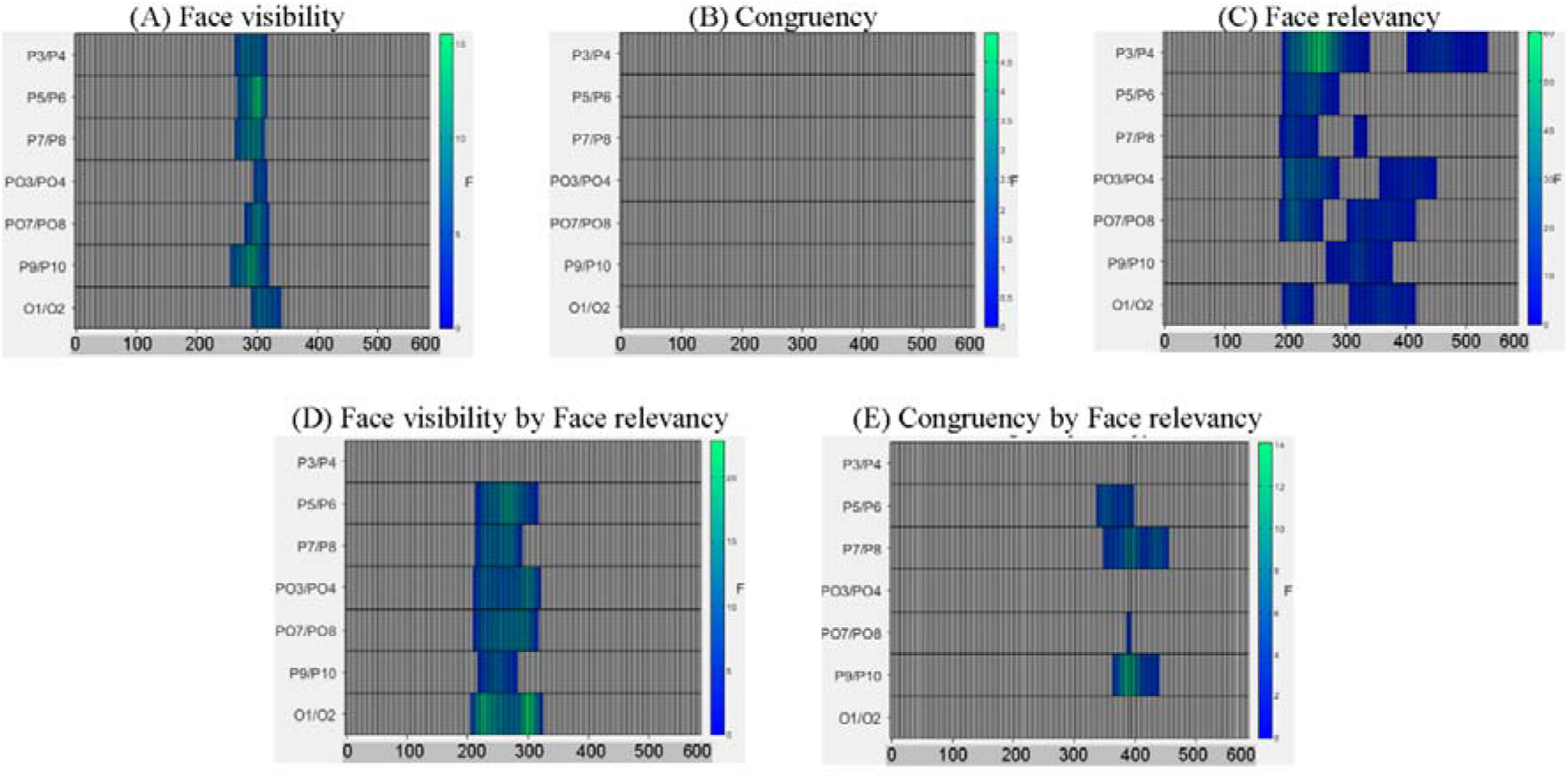
Raster plots of effects from the 3(fearful-face-dot congruency) x 2(face-visibility) x 2(face-relevancy) repeated-measures ANOVA.

To analyse the interaction effect between face-visibility and face-relevancy on the dot-related ERPs, we examined the differences between the two face-relevancy conditions (face-irrelevant minus face-relevant), separately for when the faces were presented subliminally and when presented supraliminally.

When the preceding faces were presented subliminally, there was a significant negative cluster between 246-445ms across electrodes P7/8, PO3/4, PO7/8, P9/10 and O1/2 with the maximal effect found on O1/2 (temporal peak: 316ms), see Figure 4A. Therefore, the N2 for dots in the face-relevant condition was smaller in this time window, compared to the face-irrelevant condition.

**Figure 4.**
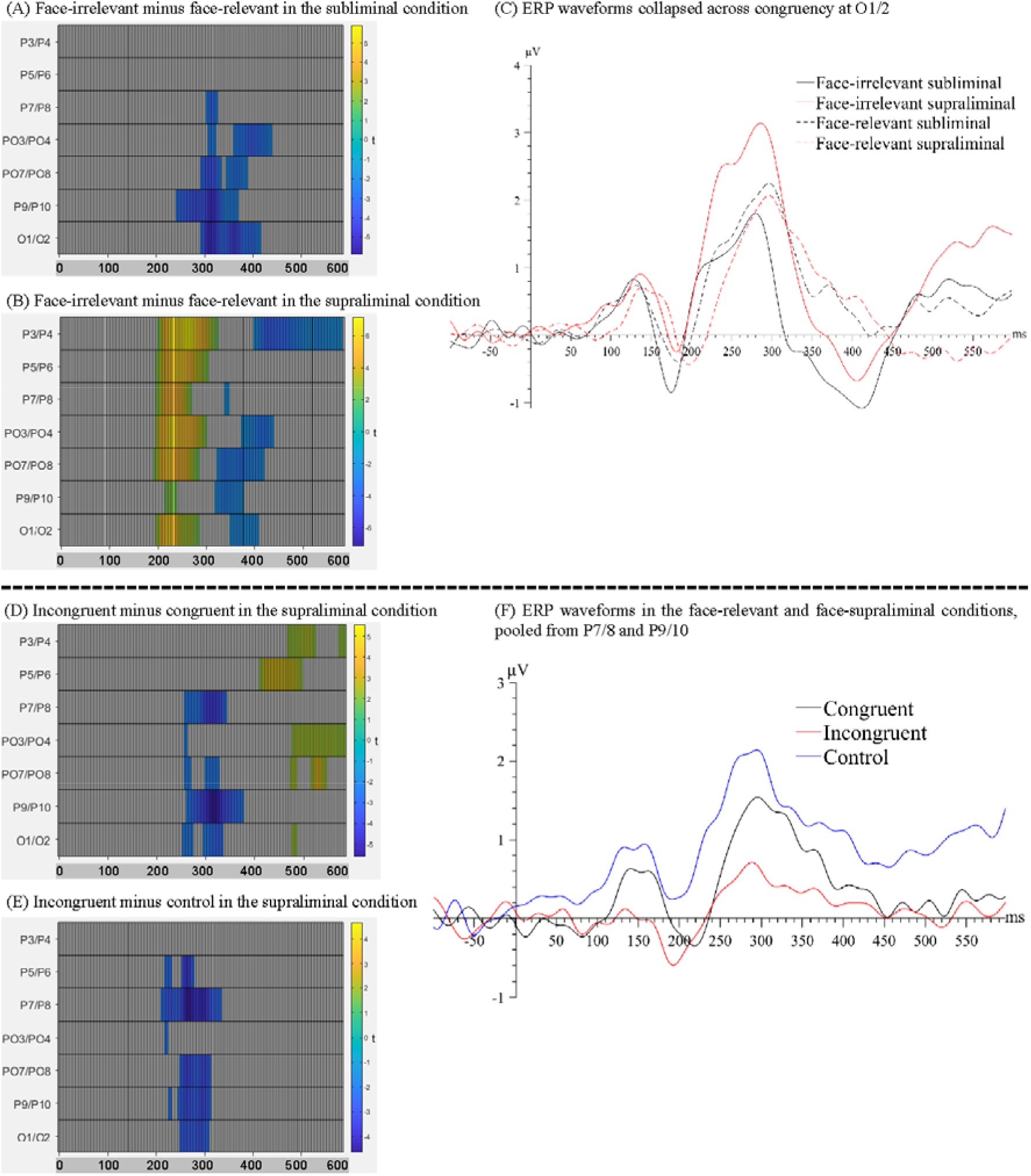
Raster plots for the contrasts between (A) face-irrelevant and face-relevant signals in the subliminal condition and between (B) face-irrelevant and face-relevant signals in the supraliminal condition. (C) ERP waveforms collapsed across congruency conditions at O1/2, the pair of electrodes showing the maximal interaction effect between face-relevancy and face-visibility. Raster plots for the contrasts between (D) incongruent and congruent signals and between (E) incongruent and control signals, in the supraliminal condition. (F) ERP waveforms for each congruency level in the face-relevant and face-supraliminal condition, pooled over P7/8 and P9/10.

When the preceding faces were presented supraliminally, there was a significant positive cluster between 195-328ms across all posterior electrodes with the maximal effect found on P3/4 (temporal peak: 266ms), see Figure 4B. Combined with the ERP waveforms (Figure 4C), it appears that the P2 for dots in the face-relevant condition was smaller than the face-irrelevant condition. In a later time window of 324-445ms, task-relevant faces seemingly continued to weaken the dot-related N2, compared to task-irrelevant faces. Therefore, the mid-latency ERPs (i.e., P2 and N2) for dots were attenuated overall by a pair of preceding task-relevant faces, especially when they were presented supraliminally (Figure 4C), potentially resulting in a slower reaction time to the target dots in the face-relevant condition.

From the omnibus analysis on dot-related signals, the main effect of congruency was not significant. However, the interaction between congruency and face-relevancy was significant (see Figure 3). A simple effect test revealed that the congruency effect was significant only in the face-relevant condition. As part of our planned comparisons, we compared levels of congruency (congruent, incongruent, control) against each other at each level of face visibility, in the face-relevant condition.

When the faces were presented subliminally, there were no significant differences in any of the comparison pairs. However, when the faces were presented supraliminally, signals in the incongruent condition were more negative than in the congruent condition between 258-383ms (spatial peak: P9/10; temporal peak: 320ms), see Figure 4D. Signals in the incongruent condition were also more negative than the control condition between 215-340ms (spatial peak: P7/8; temporal peak: 270ms), see Figure 4E. Combined with the ERP waveforms (Figure 4F), it appears that the P2 for the dots was smaller when they were presented spatially incongruent (opposite) to the preceding fearful face, compared to when they were presented at a spatially congruent location. Thus, it appears that the processing of a target dot was impaired when it was presented at an incongruent spatial location following a task-relevant and visible fearful face, and such impaired processing was accompanied by a slower reaction to the target dot.

### 3.3 Multivariate pattern analysis

#### 3.3.1 Decoding the spatial location of fearful faces

Using MVPA, we decoded the neural activity associated with the spatial locations of the fearful faces (fearful-face-on-left vs. fearful-face-on-right) in the subliminal and supraliminal conditions, separately. The decoding was successful only in the supraliminal conditions (Figure 5A, left panel), with the accuracy significantly above chance level at ∼53% (*SEM* = 0.72) between 270-289ms. However, the decoding accuracy was at chance-level in the subliminal conditions (Figure 5B, left panel). These results were in line with the ERP results in showing that the spatial location of fearful faces was decodable only in conditions where participants were aware of the face stimuli.

**Figure 5.**
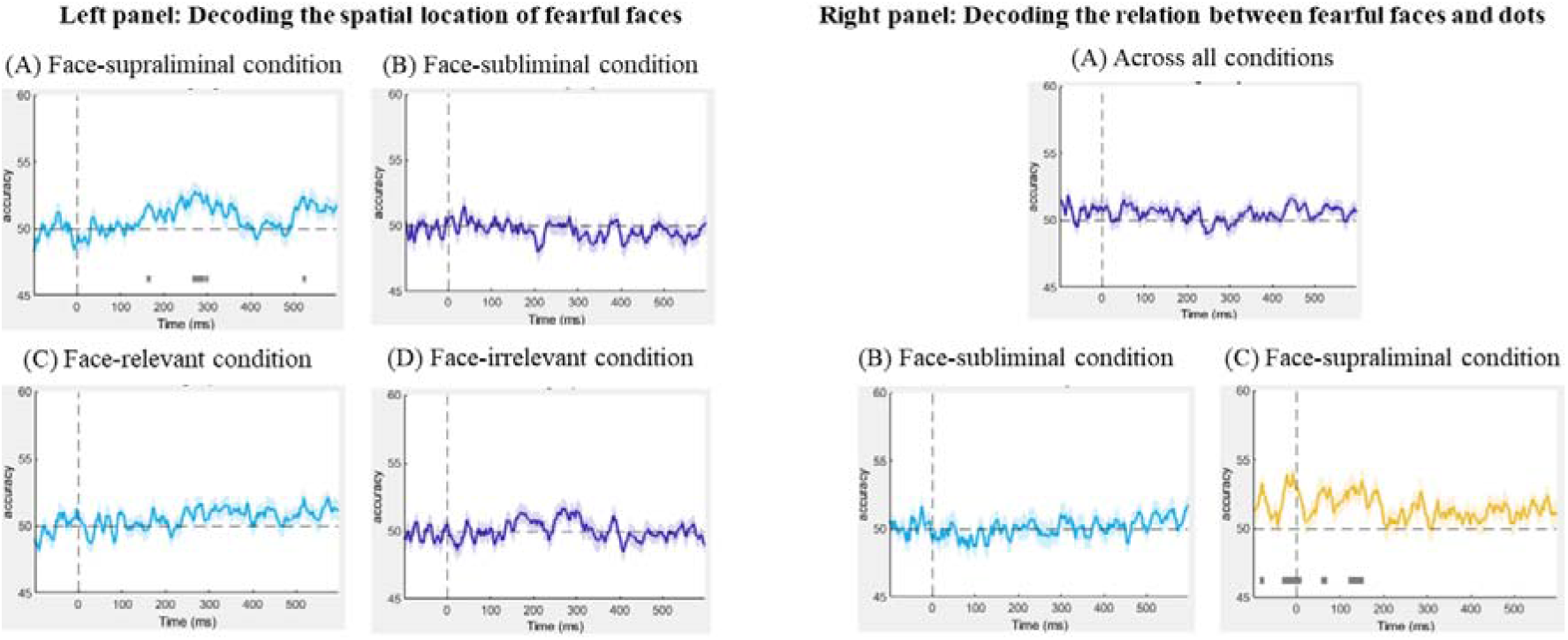
Left panel: Results of the decoding of spatial location of fearful faces in the (A) face-supraliminal condition, (B) face-subliminal condition, (C) face-relevant condition and (D) face-irrelevant condition. Right panel: Results of the decoding of the relation between fearful faces and the dots (A) across all conditions, (B) in the face-subliminal condition and (C) in the face-supraliminal condition.

We also decoded the spatial location of the fearful face separately in the task-relevant and task-irrelevant conditions, pooling over visibility. The decoding performance from both analyses was at chance-level throughout the entire epoch (Figure 5C & 5D, left panel), showing that the location of the fearful face was not decoded in either condition.

#### 3.3.2 Decoding the relation between a fearful face and the subsequent stimulus

To examine if there was any neural pattern associated with the fearful-face-dot congruency, we decoded the neural activity between fearful-face-dot congruent and fearful-face-dot incongruent trials. We performed the decoding first across all conditions and then separately for the face-subliminal and face-supraliminal conditions. No successful decoding of the fearful-face-dot relation was found overall (Figure 5A, right panel) or in the face-subliminal condition (Figure 5B, right panel). In the face-supraliminal condition, the decoding of congruency returned some significant results between −29 and 8ms (*M*_*accuracy*_ = 49%, *SEM* = 0.69) and between 123 and 150ms (*M*_*accuracy*_ = 50%, *SEM* = 0.57) (Figure 5C, right panel). Considering the near-chance decoding accuracies, we do not argue that there is very strong evidence for a successful decoding of congruency in the current analysis.

## 4. Discussion

Using the dot-probe paradigm with backward masked faces, we examined neural activity associated with the processing of fearful expressions, as well as subsequent visual targets.

From our mass univariate analysis on the dot-evoked signals, we found that, in the supraliminal face-relevant condition, mid-latency ERP signals (i.e., the P2) were stronger for dots that appeared at the same locations as the fearful face (congruent condition), compared to when they were presented at locations opposite the fearful face (incongruent condition). Behaviourally, we found evidence for this cue validity effect, again only when the preceding face stimuli (cues) were presented supraliminally and when they were task-relevant. These results are consistent with numerous reports of the facilitatory effects of an emotional face on validly cued stimuli (for a review see Torrence & Troup, 2018). Crucially, this effect was not observed when the preceding faces were presented subliminally or when they were task-irrelevant. Therefore, the fearful face-related modulatory effect on subsequent stimuli requires conscious awareness of and top-down attention to the faces. In addition, our MVPA revealed a very low accuracy for congruency decoding. One potential reason for this could be that, in the MVPA, the baseline un-subtracted signals (dot-present ERPs) were used, and more noise introduced by neural patterns associated with the preceding face stimuli was present in the data, which resulted in a rather noisy and low decoding performance of the variable of interest (congruency).

Our second main finding is that supraliminally-presented fearful faces elicited an N2pc towards them, regardless of whether the faces were task-relevant or not. While this finding is consistent to some existing research (e.g., Bar-Haim et al., 2005; Eimer & Kiss, 2007), inconsistent conclusions have been made in more recent studies including our own work (Lien et al., 2013; Qiu et al., 2022a, 2023; Zhou et al., 2020).

Although we failed to find an N2pc for task-irrelevant fearful faces in our previous experiments (Qiu et al., 2022a, 2023), our current findings do not contradict these previous findings. Specifically, task-relevancy was implemented via different methods across studies. The task-induced attentional load (Lavie, 2005; Lavie et al., 2004) varied largely between the current study and our previous ones, which could have led to two distinctly different conclusions. In our previous studies, the target non-face stimuli were superimposed onto the faces, and the onset of target stimuli was the same as the overlapping face images. Participants had to suppress the face information to accurately make a decision about the contrast-induced lines overlaid on the face images. Perhaps, the overall attentional load was higher in these previous studies, which could have prevented an N2pc from occurring (Qiu et al., 2022a, 2023). However, in the current face-irrelevant condition, the faces and the target stimuli were separated temporally by 66ms. The competition between the faces and the targets is considered lower in the current paradigm, potentially allowing some processing of the task-irrelevant faces. As a result, an N2pc was evoked by the fearful face in the pair.

Additionally, when the faces were made task-relevant, we observed a SPCN, a marker for working memory maintenance, for the fearful face. This means that, in face-relevant conditions, the target fearful face was encoded and maintained in working memory, perhaps because this was necessary to produce a correct answer in the fearful face localisation task. However, when the requirement of attending to the fearful face was removed in the face-irrelevant condition, we no longer observed the neural processes associated with working memory (SPCN). These results are not surprising as the top-down suppression of task-irrelevant signals have been demonstrated extensively in the literature (e.g., Gaspelin & Luck, 2018; Liu et al., 2020; for a review see Luck et al., 2021). As explained, the temporal separation between the two pairs of stimuli (faces and target dots) may have allowed attention to shift to task-irrelevant fearful faces. This is not incompatible with the “inhibitory mechanism” highlighted in this line of research where the inhibition was oftentimes exerted upon salient task-irrelevant stimuli presented simultaneously with the targets (Luck et al., 2021; for studies on face processing see Lien et al., 2013; Qiu et al., 2022a, 2023; Zhou et al., 2020). In the current data, signal suppression was present still, however it manifested as an absence of working memory consolidation for the task-irrelevant faces.

The task-relevancy of the faces also modulated the processing of the following target dot in the current paradigm. Specifically, a pair of task-relevant faces weakened the overall neural activity for the subsequent targets, regardless of the visibility of the faces. Perhaps, the dual task demands resulted in a higher attentional load in the face-relevant condition. Specifically, the task-relevant faces required a certain amount of the limited attentional or working memory resources (Ma et al., 2014; Xu & Chun, 2009) that are shared by neural processes for the target dots in close temporal proximity. Consequently, the strength of neural activity associated with the dots decreased as the preceding faces were processed more strongly in a task-relevant situation, compared to the face-irrelevant condition. Further, the decrease in dot-related ERPs was more evident when the faces were clearly visible (in the supraliminal condition) whereby the time range (200-400ms) of this attenuation effect was larger than when the faces were presented subliminally (300-400ms). It is likely that faces were more distracting when they were clearly visible, resulting in even stronger neural activity for the faces themselves, but further diminished ERPs for the subsequent dots. This finding is in line with our previous study using two rapid streams of visual presentations of faces, from which the amplitude of the N2pc towards a lateralised fearful face was found to decrease substantially when participants had to attend to another pair of faces presented immediately prior to it, compared to when the two face pairs were separated for longer (Qiu et al., 2022b).

Finally and most importantly, no evidence was found for the nonconscious processing of fearful faces in the current paradigm. This was supported by both the univariate and multivariate analysis results, and through two indices of spatial attention (face-related signals and dot-related signals). Indeed, no N2pc for fearful faces was observed in the subliminal viewing condition, and the neural processes for the dots following subliminal face presentations were not modulated by the presence of a fearful face. Consistent with this, the MVPA decoding performance for the spatial location of the fearful faces was at chance-level when the faces were presented subliminally. Additionally, no successful decoding of congruency was found in the face-subliminal condition. Thus, in a bilateral presentation of face images, the spatial information about a lateralised fearful face cannot be processed without visual awareness. This finding is consistent with our recent studies (Qiu et al., 2022a, 2023) and several studies by other researchers (Baier et al., 2022; Gray et al., 2013; Hedger et al., 2013; Koster et al., 2007; Tipura & Pegna, 2022).

The majority of studies showing evidence for the nonconscious processing of fearful faces used central face presentations (e.g., Del Zotto & Pegna, 2015; Pegna et al., 2008, 2011). Faces presented laterally or more eccentric in the visual field are usually harder to detect, compared to faces presented at the centre of the visual field (Papaioannou & Luck, 2020; Smith & Rossit, 2018). Even when they are presented in a subliminal viewing condition, i.e., 16ms (Del Zotto & Pegna, 2015; Pegna et al., 2008), some processing of fearful faces may occur for centrally presented faces, which may then result in a modulation of the ERPs. However, this modulation is not observed for lateralised faces, as shown by the current results, perhaps due to competition between the two similarly complex face stimuli in each presentation (Wirth & Wentura, 2018).

The use of bilateral presentations of faces in other studies on nonconscious emotion processing is however not rare (Bertini et al., 2013; Carlson & Reinke, 2008, 2010; Cecere et al., 2014; De Gelder et al., 2005; Hedger et al., 2013; Koster et al., 2007). For example, in Carlson and Reinke (2010), face pairs were presented for 33ms in a backward masking experiment. It was found that the face-sensitive N170 for the masked fearful faces was enhanced, compared to masked neutral faces. However, as acknowledged by the authors themselves, there may be some conscious experience of the stimuli when they are presented for 33ms (Carlson & Reinke, 2010; see also Pessoa et al., 2005), potentially accounting for the fear-related enhancement effect. In situations where visual awareness was more strongly impeded by either shorter presentation of faces in masking experiments (17ms, Hedger et al., 2013; 14ms, Koster et al., 2007) or in a continuous flash suppression procedure (Hedger et al., 2013), fearful faces did not attract spatial attention at the behavioural level. Our current EEG data complement the previous literature by showing that spatial attention was not captured by lateralised fearful faces presented subliminally (16ms).

Another approach to investigate nonconscious emotion processing is testing patients with cortical blindness. Such patients usually experience a loss of visual awareness due to regional lesion(s) in the brain. Previous clinical studies have consistently shown that emotional faces can be processed even though the patients were incapable of detecting or reporting the stimuli (for a review see Celeghin et al., 2015). Specifically, in a bilateral presentation of face images, patients with hemifield blindness showed improved task performance on face stimuli (e.g., better emotion recognition) presented in their intact visual field when a fearful face was concurrently presented in the blind visual field, indicating some processing of the fearful face in the absence of awareness (Bertini et al., 2013; Cecere et al., 2014; De Gelder et al., 2005). This fear-related improvement on task performance was supported by electrophysical evidence (Cecere et al., 2014) as well as functional imaging evidence (De Gelder et al., 2005). However, while fearful faces can be processed (or “influence cognitive processing”, Koster et al., 2007) nonconsciously, they do not necessarily attract spatial attention. Supporting this, in a patient with complete destruction of the primary visual cortex, Del Zotto and colleagues (2013) demonstrated that, while the presence of emotional faces, compared to neutral faces, facilitated the patient’s task performance for subsequent sound stimuli, the spatial location of the emotional faces had no effect on the patient’s behaviour (see supplementary data in Del Zotto et al., 2013).

Taken together, we conclude that, when not consciously detected, fearful faces do not attract spatial attention and they do not affect the processing of spatially contiguous stimuli. Although consciously seen fearful faces attract spatial attention and they modulate the neural processes for following stimuli, these processes are strongly modulated by attentional load. As part of the endeavour in understanding emotional face processing, the current results point to the importance of various conditions (i.e., awareness and task-relevancy) for attentional capture by fearful faces.

## Data Availability Statement

The datasets for this study can be found here: https://osf.io/54nw2/

## Author Contributions

ZQ and AJP contributed to conception and design of the study. ZQ and JJ collected data and performed data processing. ZQ performed data analyses. ZQ wrote the first draft of the manuscript. ZQ, SIB and AJP contributed to manuscript revision. All authors read and approved the submitted version.

## Acknowledgements

ZQ was supported by UQ PhD scholarships.

